# Adaptive-like NK cell responses to influenza correlate with humoral immunity and are influenced by age and sex

**DOI:** 10.1101/2025.05.24.655902

**Authors:** Eric Alves, Jared M. Oakes, Joshua D. Simmons, Jennifer Currenti, Jerome D. Coudert, Bree Foley, Joan Eason, Natasha B. Halasa, H. Keipp Talbot, Jessica L. Castilho, Simon A. Mallal, Silvana Gaudieri, Spyros A. Kalams

## Abstract

Influenza remains a global health threat, infecting approximately one billion people annually and causing significant mortality, particularly among older adults. While hemagglutination inhibition (HAI) antibody titers are a standard correlate of immunity against influenza, they do not reliably predict protection in high-risk populations. Using multiomic single-cell profiling, we identified a distinct subset of adaptive-like NK cells that respond to influenza antigen, predominantly in younger females. These *TNFSF10*^+^*LGALS9*^+^ NK cells exhibit features of adaptive NK cells but lack classical cytomegalovirus-driven markers observed in previous studies. Notably, their increased frequency correlates with high pre-existing HAI titers, suggesting a link between adaptive-like NK responses and humoral immunity. Together, our findings identify an NK subset influenced by age and sex that may contribute to influenza protection, expanding the known diversity of adaptive-like NK cells. These insights could inform future vaccine strategies, particularly for aging populations, by integrating NK responses into assessments of vaccine efficacy.

## 1. Introduction

Influenza virus continues to pose a major global health threat, infecting approximately one billion people annually, and causing an estimated 290,000 to 650,000 respiratory deaths, predominantly in adults aged 65 years and older (*1*). Despite the availability of vaccines, influenza virus undergoes continuous antigenic evolution through the accumulation of mutations (*2, 3*) or re-assortment (*4, 5*). This ongoing evolution of viral strains enables immune evasion, resulting in reinfections, even among individuals with prior natural or vaccine-induced immunity. Consequently, influenza vaccines require frequent reformulation to match circulating strains and combat antigenic evolution, yet their efficacy can vary widely between seasons and populations (*6, 7*).

Antibodies targeting influenza virus surface glycoproteins, particularly hemagglutinin (HA), have traditionally been considered the primary correlate of protection and a key measure of vaccine efficacy (*8, 9*). However, hemagglutination inhibition (HAI) antibody titers are not consistently reliable predictors of protection, especially for high-risk populations such as older adults, highlighting the need for additional immunological markers to better assess immunity (*10, 11*).

Recent analysis of the humoral immune response to influenza in a cohort of vaccinated older adults (>65 years) during an influenza season demonstrated robust antibody responses against both vaccine and circulating strains (*12*). However, individuals who later became infected exhibited distinct antibody effector profiles, marked by a deficiency in natural killer (NK) cell-activating antibodies. Notably, current recommendations for high dose vaccination in older adults failed to induce this NK cell-activating response. These findings suggest that NK cells play an important role in protection against influenza, particularly in older adults. However, the specific NK cell subset(s) responsible for this protection remain unclear.

While traditionally classified as components of the innate immune system, NK cells have increasingly been recognized for exhibiting key features of adaptive immunity (*13, 14*). To date, cytomegalovirus (CMV) infection has served as a model demonstrating that human NK cells can display antigen specificity, undergo clonal expansion, and generate stable, long-lived memory populations (*15–17*). Based on these observations, we hypothesized that the NK cell subset(s) contributing to anti-influenza immunity may represent a population of influenza-specific “adaptive” NK cells. Given the limited understanding of NK cell involvement in vaccine-induced immunity to influenza, we sought to characterize these responses at high resolution.

To investigate this, peripheral blood mononuclear cells (PBMCs) were collected from younger (<45) and older (≥65) adults seven days after seasonal influenza vaccination, coinciding with the peak of adaptive immune responses (*18*), and stimulated overnight in the presence (or absence) of inactivated H1N1 A/Brisbane/02/2018 and H3N2 A/Kansas/14/2017 - components of the vaccine administered one week earlier. We then performed multiomic single-cell profiling to conduct an unbiased, high-resolution analysis of NK cell recall responses to influenza antigen in younger and older adults. This approach enabled a comprehensive characterization of peripheral blood NK cell heterogeneity at seven days post-vaccination, comparing unstimulated cells (media control) to those stimulated with the vaccine-targeted influenza antigen, and revealed dynamic, age-specific shifts in NK cell subsets. In doing so, we identified a distinct subset of age- and sex-associated NK cells responsive to influenza antigen, exhibiting hallmarks of adaptive NK cells. Finally, we examined the relationship between these influenza antigen-reactive adaptive-like NK cells and pre-existing HAI antibody titers, providing insight into their potential link to humoral immunity.

## 2. Results

### 2.1. Single-cell profiling reveals the diverse landscape of peripheral NK cells post-vaccination

To investigate cellular responses to influenza vaccination across the adult lifespan, we generated a single-cell dataset of peripheral blood mononuclear cells (PBMCs) stimulated *in vitro* either in the presence of influenza antigen (A/Brisbane/02/2018 and A/Kansas/14/2017, matching vaccine strains administered one week prior; hereafter referred to as the “influenza antigen” condition) or in its absence (the “media” control condition). This dataset included samples from 12 younger adults (median age 28, range 23-44 years) and 12 older adults (median age 70, range 65-77 years), all of whom had received annual influenza vaccines since the 2016-2017 season, with PBMCs collected at three timepoints each season: pre-vaccination (day 0), day 7 and day 28 post-vaccination. Single-cell RNA sequencing was performed on PBMCs collected seven days post-vaccination during the 2019-2020 influenza season, a time point coinciding with peak adaptive immune responses (*18*), and likely representing an optimal window for assessing recall responses. A subset of subjects also underwent profiling for cell surface protein expression. In our companion paper, we focused on the B cell, T cell and myeloid cell compartments (*19*). Here, we specifically explore the NK cells, given emerging evidence suggesting that they may play a role in protection against influenza (*12*). Of the 411,167 total PBMCs profiled (*19*), we identified and analyzed 42,368 NK cells, derived from 11 of the 12 younger adults (excluding one younger male with no NK cells in the media control condition) and all 12 older adults (Figure 1A & 1B). Of these, 25% (10,901 cells) had paired cell surface protein data (Figure 1B).

**Figure 1.**
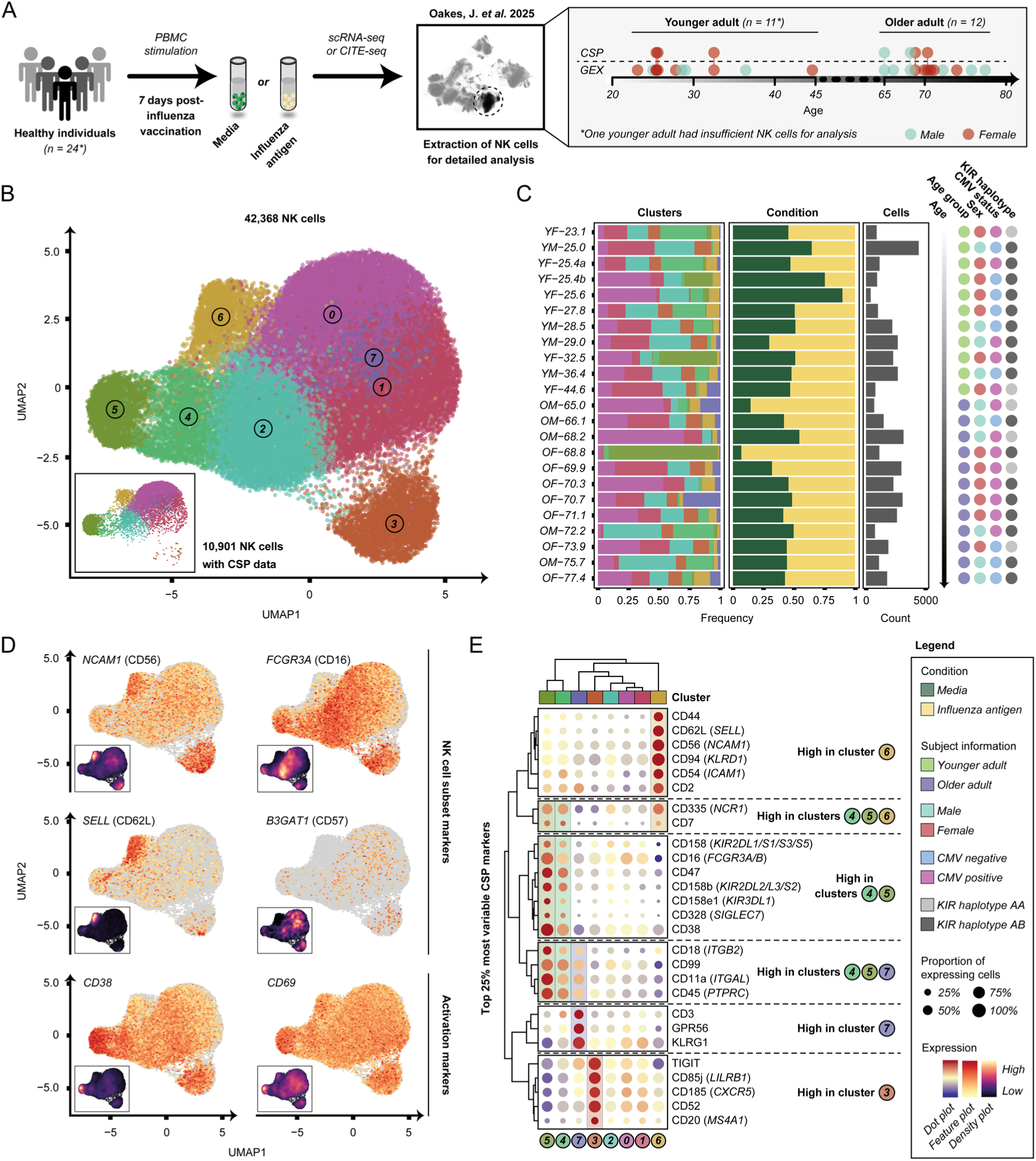
NK cells post-vaccination exhibit recall responses to influenza antigen. **A)** Schematic of the experimental design illustrating the *in vitro* stimulation of peripheral blood mononuclear cells (PBMCs) from recently vaccinated younger (*n* = 11) and older (*n* = 12) adults, either in the presence of vaccine-matched influenza antigen (yellow) or in its absence (media control; green). NK cells were extracted for detailed analysis. Single-cell gene expression (GEX) was performed for all subjects, and a subset *(n* = 3 younger adults; *n* = 4 older adults) underwent additional cell surface protein (CSP) barcoding. **B)** Uniform Manifold Approximation and Projection (UMAP) plot of 42,368 NK cells based on GEX data, with a subset (10,901 NK cells) having both GEX and CSP data. **C)** Frequency of cells in each cluster, frequency of cells profiled in either the influenza antigen or media control conditions, and total cell count profiled per subject. Subjects are ordered by age, with sex, cytomegalovirus (CMV) status and killer immunoglobulin-like receptor (KIR) haplotype indicated. Subject naming refers to younger (Y), older (O), male (M) and female (F). **D)** Feature plots and corresponding density plots showing gene expression of key NK cell subset markers (*NCAM1*, *FCGR3A*, *SELL* and *B3GAT1*) and activation markers (*CD38* and *CD69*). **E)** Dot plot displaying the 25% most variable cell surface proteins across UMAP clusters.

Unsupervised clustering identified eight NK cell populations, with seven containing cells from all subjects (Figure 1B & 1C). Cluster 5, however, was absent in three subjects. Each subject contributed a median 1,776 NK cells (range, 401-4,422 cells), spanning both influenza antigen (1,048; 41-2,019 cells) and media control (758; 78-2,857 cells) conditions (Figure 1C). Moreover, the number of NK cells captured per condition did not differ significantly (*p*-value = 0.098; Wilcoxon signed-rank test), which is consistent with prior observations (*20*), showing that overnight antigen stimulation is insufficient to induce notable NK cell expansion.

Using the transcriptional (Figure 1D) and cell surface protein (Figure 1E) signatures of each cluster, we identified cluster 6 as a CD56^bright^ NK population, characterized by high surface protein expression of CD56 and CD62L, and low levels of CD16. This subset comprised a comparable median 4.9% (range, 0.5-13.9%) of cells in the influenza antigen condition, and 6.3% (1.3-12.4%) of cells in the media control condition, consistent with expected peripheral blood frequencies (*21*).

Clusters 0, 1 and 2 were identified as CD56^dim^ NK cell populations, exhibiting low *NCAM1* and intermediate *FCGR3A* gene expression. These clusters accounted for a median 25.5% (range, 0.5-83.9%), 19.1% (1.9-40.2%) and 19.7% (1.3-39.0%) of cells in the media control condition, and 9.9% (0-53.7%), 18.2% (0-44.6%) and 14.8% (0-68.8%) of cells in the influenza antigen condition, respectively.

Cluster 7, comprising a median 2.9% (range, 0-30.5%) of cells in the media control and 2.5% (0-30.7%) of cells in the influenza antigen conditions, displayed high cell surface protein expression of GPR56, KLRG1, and unexpectedly, CD3. However, these cells did not express *CD3D* (adjusted *p*-value = 1; average log_2_FC = 1.08), *CD3E* (adjusted *p*-value = 1; average log_2_FC = 1.01) or *CD3G* (adjusted *p*-value = 1; average log_2_FC = 1.04) transcripts at levels above those of other NK clusters, suggesting the surface CD3 signal may arise from trogocytosis following interactions with T cells (*22, 23*). This interpretation is supported by the culture conditions, in which NK cells were incubated overnight alongside T cells and other PBMCs prior to single-cell sequencing.

Clusters 4 and 5 both exhibited a similar CD56^dim^ phenotype, but were distinguished from clusters 0, 1 and 2 by high gene expression of *killer immunoglobulin-like receptors (KIRs)*, *CD38* and *CD69*, indicative of activation. Cluster 4 represented a median 3.0% (range, 0.1-60.1%) of cells in the media control and 3.3% (0-65.1%) of cells in the influenza antigen conditions, whereas cluster 5 was comparatively rare (median 0.1% [range, 0-4.5%] in media control; 0.6% [0-95.8%] in influenza antigen).

Finally, cluster 3 exhibited a distinct cell surface protein signature, marked by high surface protein levels of TIGIT, CD85j, CD185, CD52 and CD20, alongside moderate *NCAM1* and *FCGR3A* gene expression. This cluster accounted for a median 5.8% (range, 0-19.6%) of cells in the media control and 5.9% (0-18.6%) of cells in the influenza antigen conditions.

Collectively, these findings reveal substantial NK cell heterogeneity in peripheral blood post-vaccination, with all major subsets present across subjects. However, variation in the relative abundances of specific clusters across subjects suggests potential age-related influences on NK cell composition.

### 2.2. Influenza antigen stimulation alters NK cell subset frequencies in younger but not older adults post-vaccination

We next examined the distribution of post-vaccination NK cell subsets in the presence or absence of vaccine-matched influenza antigen, stratified by age. Unsupervised clustering revealed differences in cluster frequencies across age groups and stimulation conditions (Figure 2A). In the media control condition, cluster 4 was the only cluster that differed significantly between younger and older adults, being higher in older adults (FDR = 0.031). The remaining clusters showed no significant age-related differences in abundance (cluster 0: FDR = 0.075; cluster 1: FDR = 0.74; cluster 2: FDR = 0.52; cluster 3: FDR = 0.88; cluster 5: FDR = 0.084; cluster 6: FDR = 0.97; cluster 7: FDR = 0.084).

**Figure 2.**
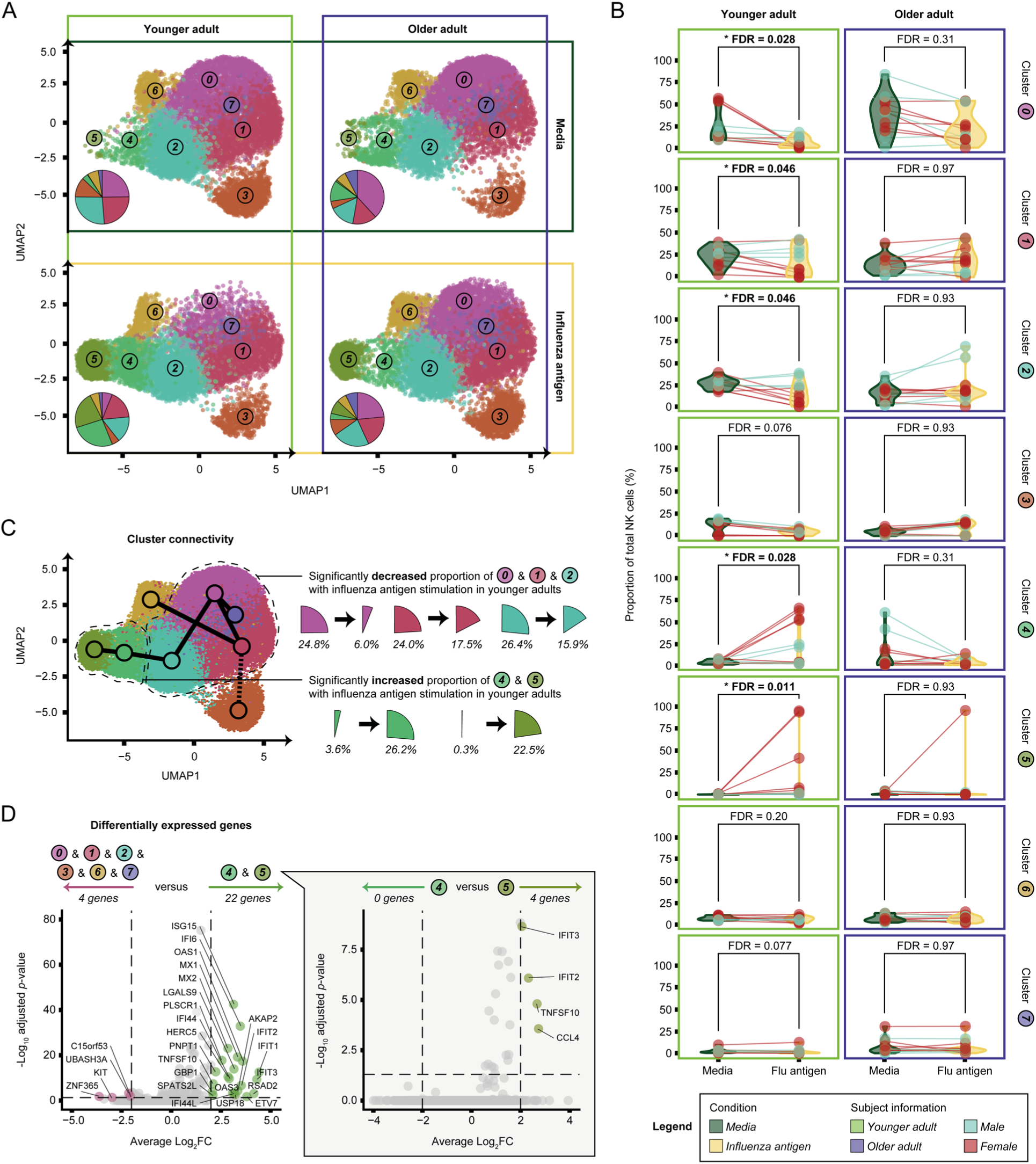
Influenza antigen modifies the distribution of post-vaccination NK cell subsets. **A)** UMAP plots stratified by age group and stimulation condition. Pie charts illustrate the distribution of NK cell populations across clusters. **B)** Changes in cluster frequency between media control and influenza antigen conditions, shown separately for each age group. Significant changes (FDR < 0.05) are indicated in bold. Analyses conducted using paired FDR-corrected moderated t-tests. **C)** Cluster connectivity inferred from lineage trajectory analysis. Solid lines represent stable connections, while the dotted line indicates a connection classified as an outlier. The findings from (B) are visually summarized. **D)** Differential gene expression analysis comparing clusters 4 and 5 (which show a significant frequency increase in the presence of influenza antigen) to all other clusters combined. A separate comparison of cluster 4 versus cluster 5 is also shown. Differentially expressed genes are labelled, with horizontal and vertical black dotted lines indicating statistical thresholds for significance (average Log_2_FC > |2| and adjusted *p*-value < 0.05).

In the presence of influenza antigen, younger adults exhibited significant shifts in NK cell subset frequencies, whereas older adults showed no consistent changes (Figure 2B). In younger adults, clusters 4 (FDR = 0.028) and 5 (FDR = 0.011) were significantly enriched in the influenza antigen condition, with cluster 5 observed almost exclusively in this condition (only one older adult, OF-68.8, showed a high frequency of cluster 5 NK cells). In contrast, clusters 0 (FDR = 0.028), 1 (FDR = 0.046) and 2 (FDR = 0.046) were more prevalent in the media control condition. The frequencies of clusters 3 (FDR = 0.076), 6 (FDR = 0.20) and 7 (FDR = 0.077) remained largely unchanged. Notably, an inverse trend in cluster 3 frequency with age was observed, though it did not reach statistical significance (Figure S1). Together, these findings suggest that younger adults mount a more dynamic NK cell response to influenza antigen stimulation, whereas older adults exhibit a more static NK cell profile.

To better understand the shifts observed in younger adults, we performed lineage trajectory analysis to explore cluster connectivity (Figure 2C), which suggested that clusters 0, 1, 2, 4 and 5 may share a lineage. This analysis supports the observation that these clusters are the most affected by influenza antigen stimulation, and also suggests that cluster 5 – an activated subset largely absent in the media control condition – may arise from resting cells within clusters 0, 1 and/or 2.

Given the enrichment of activated NK cell clusters 4 and 5 following influenza antigen stimulation, we next characterized their transcriptional profiles. Differential gene expression analysis comparing clusters 4 and 5 (combined) to all other NK cell clusters, regardless of condition, identified 22 significantly upregulated genes (Figure 2D; Table S1). These included multiple interferon-stimulated genes (*IFIT1*/*2*/*3*, *IFI6*/*44*/*44L*, *RSAD2*, *OAS1*/*3*, *ISG15*, *HERC5*, *GBP1*, *MX1*/*2* and *ETV7*), which are strongly associated with anti-viral responses. Notably, key markers of NK cell activation, including *TNFSF10*/TRAIL (adjusted *p*-value < 0.001; average log_2_FC = 2.11) and *LGALS9*/Galectin-9 (adjusted *p*-value < 0.001; average log_2_FC = 3.16), were also significantly upregulated. Additionally, we observed significant downregulation of four genes, including *KIT* (adjusted *p*-value = 0.044; average log_2_FC = 2.97), a marker typically associated with CD56^bright^ NK cells (*24*). A comparison of clusters 4 and 5 to other CD56^dim^ NK cells (clusters 0, 1, and 2) revealed a similar enrichment of activation and interferon-stimulated genes (Table S2), reinforcing the hypothesis that clusters 4 and 5 represent activated CD56^dim^ NK cell subsets.

To further refine the distinction between clusters 4 and 5, we performed differential gene expression testing between the two clusters (Figure 2D; Table S3). While no genes were uniquely enriched in cluster 4 relative to cluster 5, we identified four genes that were significantly upregulated in cluster 5: *CCL4* (adjusted *p*-value < 0.001; average log_2_FC = 2.73), *TNFSF10*/TRAIL (adjusted *p*-value < 0.001; average log_2_FC = 2.67), *IFIT2* (adjusted *p*-value < 0.001; average log_2_FC = 2.32) and *IFIT3* (adjusted *p*-value < 0.001; average log_2_FC = 2.06). These findings indicate that cluster 5 represents a more highly activated subset compared to cluster 4, further supporting the conclusion that younger adults are more likely to exhibit a more pronounced NK cell response to influenza antigen stimulation.

### 2.3. Adaptive-like NK cells are responsive to influenza antigen

Given that the subjects in this cohort are repeat influenza vaccinees with multiple prior exposures to influenza antigen, we hypothesized that clusters 4 and 5 may represent “adaptive-like” NK cells engaged in an influenza-specific recall response. While adaptive NK cells have been well characterized in the context of CMV infection – typically defined by NKG2C expression (*16, 17, 25–27*) – their role in the anti-influenza response remains poorly understood. Notably, studies of Epstein-Barr virus (EBV) infection have demonstrated that NKG2C^+^ adaptive NK cells are CMV-specific, rather than EBV-specific (*28*), suggesting that this marker may not reliably identify other virus-specific memory NK cells. To address this limitation, a broader “adaptive-like” phenotypic signature has been proposed by others (*29*), characterized by high surface expression of CD16, CD57, CD271, CD2, CD18, CD49d and killer immunoglobulin-like receptors (KIRs). To determine whether clusters 4 and 5 align with this adaptive-like NK cell profile, we assessed the expression of these markers across all NK cell clusters (Figure 3A). Although CD271 was absent from our cell surface protein panel, both clusters 4 and 5 exhibited features consistent with an adaptive-like NK cell phenotype, with cluster 5 displaying a more pronounced adaptive signature in line with its transcriptional profile. Unlike CMV-driven adaptive NK cells, we observed no significant enrichment of *KLRC2* expression within these populations (adjusted *p*-value = 1; average log_2_FC = 1.01), suggesting that NKG2C may not be a defining feature of influenza antigen-reactive adaptive-like NK cells.

**Figure 3.**
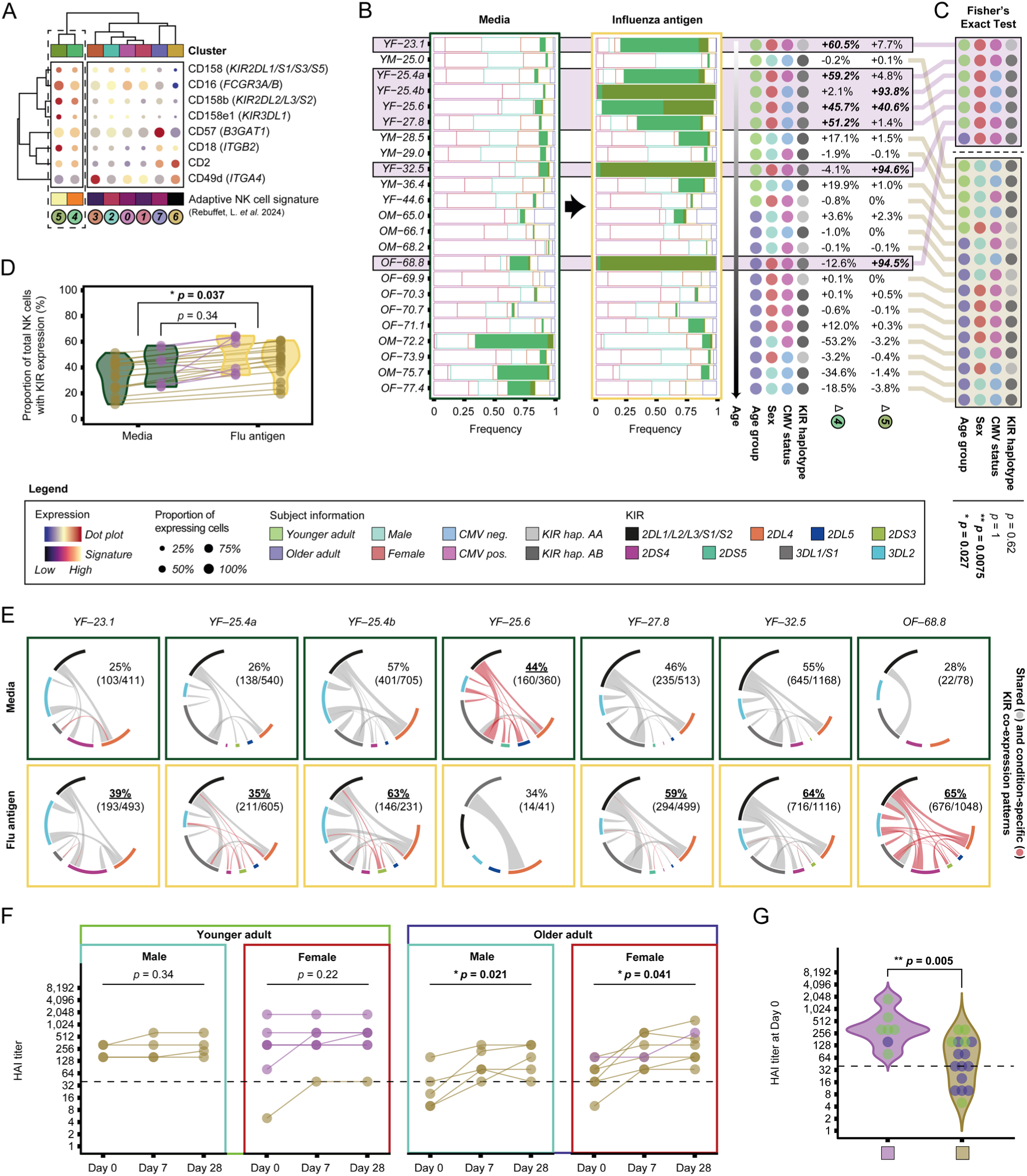
Influenza antigen-reactive adaptive-like NK cells are enriched in younger females. **A)** Dot plot of adaptive-like NK cell surface protein markers and a combined adaptive-like NK cell surface protein signature. **B)** Frequency of cells in each cluster by subject, stratified by condition. Adaptive-like NK cell clusters (clusters 4 and 5) are highlighted with filled coloring. Subjects are ordered by age, with sex, cytomegalovirus (CMV) status and killer immunoglobulin-like receptor (KIR) haplotype indicated. Frequency changes in clusters 4 and 5 in the presence versus absence of influenza antigen are shown. Major increases (>50%) in cluster 4 and/or 5 cell frequency in the presence of influenza antigen are indicated in bold. Subjects with major increases are indicated in purple. **C)** Fisher’s Exact Test of subject characteristics stratified by whether they exhibit major increases in cluster 4 and/or 5, as determined in (B). Significant characteristics (*p*-value < 0.05) are indicated in bold. **D)** Frequency of NK cells expressing one or more KIR genes, stratified by condition. All subjects combined (*n* = 23) and only subjects with major increases in clusters 4/5 (*n* = 7) were compared by paired moderated t-test. **E)** Circos plots of KIR co-expression pairs for subjects showing major increases in clusters 4/5. Co-expression pairs are only plotted if present in two or more cells. Ribbon width is indicative of co-expression pair frequency. Red ribbons indicate those present only in one condition (i.e. either media control or influenza antigen). Absence of a ribbon indicates single-KIR expression. Values indicate the proportion of NK cells expressing one or more KIRs. **F)** Hemagglutination inhibition (HAI) antibody titers at day 0 (baseline), day 7 and day 28 post-vaccination for examined subjects, stratified by age group and sex. Subjects exhibiting major increases in clusters 4/5, as determined in (B), are denoted in purple. Changes in HAI titers between time points were analyzed using repeated-measures ANOVA, with significant differences (*p*-value < 0.05) indicated in bold. **G)** Baseline HAI titers for subjects with and without major increases in clusters 4/5, as determined in (B). Comparison conducted using Mann-Whitney U Test.

We next assessed inter-individual variability in the reactivity of adaptive-like NK cell clusters in the presence of influenza antigen. Seven subjects exhibited a major (>50%) frequency increase in clusters 4 and/or 5, compared to media control (Figure 3B). Notably, all were female (*p*-value = 0.0075), and all but one were younger adults (*p*-value = 0.027), suggesting that sex and age may influence the responsiveness of influenza antigen-reactive adaptive-like NK cells (Figure 3C). Moreover, this reactivity was not restricted to CMV-positive subjects (*p* = 1), further supporting that these influenza antigen-reactive adaptive-like NK cells are unlikely to be CMV-related.

Given the role of KIRs in the adaptive NK cell response to CMV infection (*30*), we next explored whether subjects exhibiting high frequencies of influenza antigen-reactive adaptive-like NK cells share carriage of one or more specific KIR genes. KIRs serve to regulate NK cell effector function through a balance of inhibitory and activating signals, and while their expression is stochastic in healthy individuals, infection can drive preferential expansion of specific KIR-expressing subsets (*30–33*). Among the seven subjects with major increases in clusters 4 and/or 5 in the presence of influenza antigen, we observed no consistent KIR haplotype, nor did these subjects uniquely carry any specific KIR gene compared to the remaining 16 subjects (Figure S2; Table S4 & S5). The only genes shared by all seven subjects – KIR2DL4, KIR2DS4, KIR3DL2 and KIR3DL3 – were also present in nearly all other subjects (the only exception being KIR2DS4, which was absent in YM-29.0).

Next, leveraging transcriptome alignment to a KIR-modified GRCh38 genome reference (*34*), we assessed the transcriptional expression of KIR genes among the subjects. Due to the high sequence homology among KIR2DL1/L2/L3/S1/S2 and KIR3DL1/S1 – particularly at exons 1 and 2 – our short-read sequencing approach, combined with the 5’ transcript bias of the single-cell chemistry, limited our ability to accurately distinguish these genes. Nonetheless, we captured the individual expression of many KIR genes, including those absent from the cell surface protein panel. While the frequency of cells expressing *KIR2DL4*, *KIR2DS4*, *KIR3DL2* and *KIR3DL3* did not differ significantly between media control and influenza antigen conditions (Figure S2), we observed that KIR-expressing NK cells were more abundant (*p*-value = 0.037) in the influenza antigen condition compared to the media control condition, for all but one subject (YF-25.6). This elevated abundance occurred irrespective of clusters 4 and 5 (Table S6 & S7). Notably, although KIR2DL4 has been reported to be ubiquitously expressed across NK cells (*35*), we detected *KIR* transcripts in only a subset of cells – median 37.1% (range, 11.5-56.9%) in the media condition and 48.2% (19.5-64.5%) in the influenza antigen condition (Figure 3D; Table S7). This discrepancy likely reflects the well-documented challenge of transcript “dropout” in single-cell data, where genes expressed in some cells go undetected in others due to low mRNA abundance, capture efficiency and/or stochastic gene expression (*36*). Still, the consistent increase in KIR-expressing cells in the influenza antigen condition suggests a broader role for KIR regulation in the anti-influenza response.

Last, we examined whether influenza antigen stimulation altered patterns of KIR co-expression in the seven subjects with major expansions in clusters 4 and/or 5. While condition-specific co-expression patterns were evident, they were highly varied across subjects, and no consistent KIR co-expression signature emerged in the influenza antigen condition (Figure 3E; Table S6 & S7). Altogether, these findings support a role for KIRs in shaping NK cell responses to influenza antigen, though the specific patterns appear subject-dependent and heterogeneous.

### 2.4. Influenza antigen-reactive adaptive-like NK cell responsiveness associates with elevated baseline influenza antibody titers

Given the well-established role of HAI antibody titers as a primary correlate of protection and a key measure of influenza vaccine efficacy (*8, 9*), we assessed HAI titers at baseline (day 0), as well as 7- and 28-days post-vaccination (Figure 3F). Among younger adults, HAI titers remained stable over time in both males (*p*-value = 0.34) and females (*p*-value = 0.22). In contrast, older adults exhibited significant changes in HAI titers in response to vaccination (males, *p*-value = 0.021; females, *p*-value = 0.041). However, irrespective of these temporal fluctuations, younger adults had significantly higher baseline HAI titers (median, 320; range, 5-1810) compared to older adults (40; 10-160) (*p*-value = 0.004; Mann-Whitney U Test). Notably, baseline HAI titers did not significantly differ between younger males and females (*p*-value = 0.62) or between older males and females (*p*-value = 0.25). These findings reinforce the notion that younger adults tend to have a more robust pre-existing immunity to influenza, while older adults exhibit an age-related decline in antibody responses (*37*).

Strikingly, all seven subjects (7/7 female; 6/7 younger adults) with increased frequencies of adaptive-like NK cells in the presence of influenza antigen had protective (>40) baseline HAI titers (median, 320; range, 80-1810) that remained above the protective threshold at 7- and 28-days post-vaccination (Figure 3F). Compared to the remaining 16 subjects, these seven subjects had significantly higher baseline HAI titers (*p*-value = 0.005), indicative of strong pre-existing anti-influenza humoral immunity (Figure 3G). Although age may be a confounding factor, these findings raise the possibility that influenza antigen-reactive adaptive-like NK cells may influence - or be influenced by - the antibody response to influenza.

## 3. Discussion

Identifying robust correlates of protection against influenza remains a global health priority, particularly for high-risk populations, such as older adults (*10, 11*). While NK cells are recognized as key mediators of anti-viral immunity, their precise role in influenza protection – especially in aging individuals – remains incompletely understood (*12*). Here, we performed single-cell profiling of PBMCs collected seven days after seasonal influenza vaccination and stimulated *in vitro* with or without vaccine-matched influenza antigen. This approach enabled high-resolution characterization of post-vaccination NK cell responses and subset dynamics in younger and older adults.

We identified a distinct subset of *TNFSF10*^+^ (encoding TRAIL) *LGALS9*^+^ (encoding Galectin-9) NK cells that increased in frequency in the presence of influenza antigen, predominantly in younger females. These cells exhibited hallmarks of adaptive NK cells, yet they lacked key features of CMV-driven adaptive NK cells, such as upregulation of *KLRC2* (encoding NKG2C) and KIR repertoire skewing (*16, 17, 27, 30*). This suggests they represent a previously uncharacterized influenza antigen-reactive adaptive-like NK cell subset, distinct from CMV-driven populations. Notably, subjects exhibiting elevated frequencies of these cells had high pre-existing HAI titers, suggesting there may be a relationship between these influenza antigen-reactive adaptive-like NK cells and humoral immunity.

The mechanisms underlying higher frequencies of influenza antigen-reactive adaptive-like NK cells in females remain unclear. One possible factor is the influence of sex hormones. Testosterone suppresses both humoral and cellular immunity (*38*), reducing antibody responses to influenza vaccination (*39*). In contrast, estrogen enhances B cell antibody production (*40, 41*), and boosts anti-viral immune responses. Several interferon-stimulated genes, including *OAS1* and *OAS3*, which were enriched in these NK cells, exhibit female-biased expression via estrogen response elements (*42*).

Beyond hormonal influences, sex chromosome-linked mechanisms may also contribute to increased influenza antigen-reactive adaptive-like NK cells in females. Although males generally have higher circulating NK cell numbers (*43*), their NK cells exhibit reduced effector function – a disparity influenced by both sex hormones and X chromosome dosage effects (*44*). While X chromosome inactivation serves to balance gene expression between sexes, this process is incomplete. Approximately 25% of X-linked genes escape inactivation, enabling expression from both the active and inactive X chromosomes (*45*). This escape from X-inactivation has been linked to sex differences in NK cell function (*44*), and may underlie the observed sex-based differences in influenza antigen-reactive adaptive-like NK cells, warranting further investigation.

Recent work suggests that effective influenza control in both younger and older adults is associated with NK cell-activating antibodies (*12*). However, the precise NK cell subsets that mediate protection remain unknown. High-dose influenza vaccination in older adults failed to induce these NK cell-activating signals (*12*), underscoring a critical gap in our understanding of how these responses can be enhanced. Our findings build on this work by identifying an influenza antigen-reactive subset of adaptive-like NK cells that associates with humoral immunity to influenza. Notably, although sex was not explicitly analyzed in the prior study (*12*), most subjects were female (younger, 70%; older, 68%), aligning with our observation that influenza antigen-reactive adaptive-like NK cells may be influenced by sex. While further validation in larger cohorts is necessary, these findings suggest that influenza antigen-reactive adaptive-like NK cells may be a key target of NK cell-activating signals, meriting deeper investigation into their role in influenza immunity and vaccine responsiveness.

## Limitations of the study

Although the study cohort was approximately balanced by sex and age, the modest sample size poses a limitation. Nonetheless, the use of a well-characterized longitudinal cohort with repeated influenza vaccination strengthens the robustness and relevance of our findings.

A second limitation lies in the use of mixed PBMC cultures for antigen stimulation, which makes it difficult to determine whether NK cell activation was antigen-specific or occurred indirectly through bystander interactions. While future studies using isolated NK cells or defined co-cultures will be needed to resolve these mechanisms, the mixed-cell environment used here has provided key insight and may better approximate *in vivo* conditions, potentially capturing physiologically relevant interactions not observed in reductionist systems.

We also note that two subjects (YF-25.6 and OF-68.8) had fewer than 100 NK cells in one of the conditions – below the typical threshold for robust analysis (*46*) – limiting comparisons of KIR co-expression within individuals. In the context of KIR, transcript dropout and the grouping of KIR genes due to sequence homology restricted our ability to resolve a subset of the co-expression patterns across the dataset. Moreover, the overnight stimulation with influenza antigen may have been insufficient to activate specific KIR-expressing NK cell subsets. Still, despite these constraints, the KIR analysis approach and expression trends identified here provide a foundation for future analyses examining KIR using multiomic approaches.

Finally, though we observed associations between adaptive-like NK cell reactivity and pre-existing HAI titers, functional validation – such as NK cell-mediated antibody-dependent cytotoxicity (ADCC) assays – was not performed. Similarly, while trajectory analysis offered insight into the potential origin of influenza antigen-reactive adaptive-like NK cells, our computational approach does not replace direct lineage tracing experiments. Future work incorporating both functional and clonal tracking assays will be helpful to determine whether these cells directly contribute to protective immunity and to clarify their temporal dynamics.

## 4. Resource availability

### 4.1. Lead contact

Further information and requests for resources and reagents should be directed to and will be fulfilled by the lead contact, Spyros A. Kalams.

### 4.2. Materials availability

This study did not generate new unique reagents.

### 4.3. Data and code availability

- Data availability: Single-cell data will be made available upon acceptance.
- Code availability: This paper does not report original code.
- Any additional information required to reanalyze the data reported in this paper is available from the lead contact upon request.

## 5. Acknowledgements

This work was supported by a National Institutes of Health grant (R01 AI142095) awarded to SAK. EA is a recipient of an Australian Government Research Training Program Scholarship, BioZone Completion Scholarship and Fulbright Future Scholarship (funded by The Kinghorn Foundation). The funders had no role in study design, data collection and interpretation, or the decision to submit the work for publication.

We thank James F. Read and Anthony Bosco (The BIO5 Institute, University of Arizona) for their helpful recommendations on quality control of single-cell data. We also extend thanks to Charles Herring (Harry Perkins Institute of Medical Research) for his advice on single-cell data integration using downsampling.

## 6. Author contributions

Conceptualization: EA, SG, SAK.

Software: EA, JS, JC.

Formal analysis: EA, JO, JS.

Data curation: EA, JO, JS, JE, NBH, HKT, JLC, SAK.

Visualization: EA.

Supervision: SAM, SG, SAK.

Funding acquisition: SAK.

Writing – Original Draft: EA.

Writing – Review & Editing: EA, JO, JC, JDC, BF, SG, SAK.

## 8. Supplementary information titles and legends

*AlvesE_NK_SupplementaryFigures.docx*

- **Figure S1.** Cluster 3 shows opposing trends with age but lacks statistical significance.
- **Figure S2.** KIR gene-ligand repertoire does not associate with influenza antigen-associated NK cell reactivity.
- **Figure S3.** KIR allele sequence homology across the cohort.
- **Figure S4.** Identification of mismapped KIR reads.

*AlvesE_NK_SupplementaryTables.xlsx*

- **Table S1.** Genes enriched in clusters 4 and 5 compared to all other clusters.
- **Table S2.** Genes enriched in clusters 4 and 5 compared to clusters 0, 1 and 2.
- **Table S3.** Genes differentiating cluster 5 from cluster 4.
- **Table S4.** Subject HLA and KIR genotype.
- **Table S5.** Subject-specific KIR genes and corresponding ligands.
- **Table S6.** KIR expression patterns by cluster.
- **Table S7.** Cells expressing specific KIR genes by subject.
- **Table S8.** Misassignment of KIR transcripts due to allele-reference sequence homology

## 9. Experimental model and study details

Younger adults were recruited from an established cohort of healthcare workers at Vanderbilt University Medical Center, while older adults were enrolled through a Phase 4 clinical trial (NCT04077424) designed to investigate age-related differences in immune responses to influenza vaccination. All participants had received annual vaccinations during the 2016-2017, 2017-2018 and 2018-2019 seasons. For this study, samples were collected during the 2019-2020 northern hemisphere influenza season at three time points: pre-vaccination (day 0), 7 days and 28 days post-vaccination. Younger adults received a quadrivalent Fluzone (standard-dose) vaccine, while older adults received either a trivalent Fluzone High-Dose, Fluad or Flublok vaccine. Single-cell assays were performed on PBMCs isolated seven days post-vaccination.

Subjects were selected based on age and balanced as closely as possible for sex assigned at birth (referred to as “sex” throughout this study). Younger adults were predominantly female (64%; 7/11) and mostly CMV negative (55%; 6/11), while older adults had an even distribution of male and female subjects (50%; 6/12) and were primarily CMV positive (67%; 8/12). In terms of KIR haplotypes, the majority of younger (82%; 9/11) and older (67%; 8/12) adults were classified as haplotype AB, with the remainder being haplotype AA.

## 10. Method details

### 10.1. Ethics statement

All subjects gave written and verbal informed consent prior to participation and samples were anonymized. Institutional review board (IRB) approval for sample collection was obtained prior to the commencement of the study by the Vanderbilt IRB (#161647, #191639).

### 10.2. HLA and KIR genotyping

High-resolution HLA and KIR genotyping was performed at the Institute for Immunology and Infectious Diseases (IIID), Murdoch University (Perth, WA, Australia), as previously described (*47–50*). Briefly, genomic DNA was extracted from plasma using the DNeasy Blood & Tissue Kit (QIAGEN, NRW, Germany), per the manufacturer’s instructions. Locus-specific PCR amplification of genomic DNA was then performed, and amplified products were pooled in equimolar ratios for sequencing on the Illumina MiSeq platform using a 2x300 bp paired-end chemistry kit (Illumina, CA, USA). Sequencing data was quality-filtered, demultiplexed, merged and aligned using CLCbio Genomics Workbench (QIAGEN Bioinformatics). Allele calls were performed using the latest IPD-IMGT/HLA and IPD-KIR allele databases (*51*). HLA and KIR genotyping results for each subject are available in Table S4.

### 10.3. CMV serology

CMV serology was performed using the CMV IgG Enzyme Immunoassay Test Kit (GWB-892399) from Aviva Systems Biology (San Diego, CA, USA). Briefly, pre-vaccination sera were heat inactivated at 56°C for 45 minutes, followed by dilution at 1:20, 1:40 or 1:100 and incubated, along with CMV IgG standards, in duplicate on CMV antigen-coated plates at 37°C for 30 minutes. Following this, plates were washed five times with sterile PBST buffer (PBS with 0.05% TWEEN 20) and horseradish peroxidase was added for one hour at room temperature. Plates were then washed again five times with sterile PBST buffer and developed with tetramethylbenzidine substrate for 15 minutes at 37°C. Finally, stop solution (1N HCl) was added and absorbance was read at 450nm within 15 minutes.

### 10.4. HAI assay

HAI titers for each subject were measured at baseline (day 0), as well as 7- and 28-days post-vaccination, as described in our companion paper (*19*). Briefly, sera were treated with receptor-destroying enzyme (RDE; Denka Seiken, Tokyo, Japan) at 37°C for 18 hours and heat-inactivated at 56°C for 45 minutes. RDE-treated sera were diluted 1:10 in PBS. Washed turkey erythrocytes (Lampire Biological Laboratories, PA, USA) were resuspended and incubated with diluted serum at 4°C for one hour with periodic inversion. Following centrifugation, the supernatant was extracted. Viral antigens (A/Brisbane/02/2018, A/Kansas/14/2017, B/Colorado/06/2017, B/Phuket/3073/2013) were standardized to 4 HA units/25uL. The treated serum were serially diluted 2-fold in duplicate, mixed with an equal volume of antigen and incubated at room temperature for 30 minutes. Subsequently, 50uL of 0.5% turkey erythrocytes were added and incubated at room temperature for 30-45 minutes. The HAI titer was recorded as the highest dilution preventing hemagglutination.

### 10.5. PBMC stimulation

PBMCs from seven days post-vaccination were stimulated with inactivated influenza virus A/Brisbane/02/2018 and A/Kansas/14/2017, as described in our companion paper (*19*). Briefly, PBMCs were incubated with 0.5ug/mL of anti-CD40 blocking antibody (Miltenyi Biotec, CA, USA) at 37°C for 15 minutes. Following this, 1.5-2 million blocked cells were incubated overnight at 37°C with either media control (RPMI-1640, 10% human serum, 1X penicillin-streptomycin, 10mM HEPES buffer and 2mM L-glutamine) or influenza antigen-containing media (normalized to 16 HA units). Following overnight stimulation, PBMCs were washed twice with Cell Staining Buffer (BioLegend, CA, USA) and counted for pooling prior to cell surface protein labelling or single-cell sequencing.

### 10.6. Cell surface protein labelling

For a subset of subjects (*n* = 3 younger adults; *n* = 4 older adults), media control and influenza antigen-stimulated PBMCs were labelled with cell surface protein barcodes, as previously described (*52*). Briefly, cells incubated with Human TruStain FcX (Fc Receptor Blocking Solution) (BioLegend, CA, USA) at 4°C for 10 minutes, and subsequently stained with the TotalSeq-C Human Universal Cocktail V1.0 (BioLegend, CA, USA) at 4°C for 30 minutes, which labels 130 unique cell surface antigens and 7 isotype controls. Stained cells were then washed twice with Cell Staining Buffer and counted for pooling prior to single-cell sequencing.

### 10.7. Single-cell sequencing

Media control and influenza antigen condition PBMCs were pooled equally such that two samples were multiplexed per single-cell reaction. Each reaction always contained at least one media and one influenza antigen-stimulated sample, originating from one younger and one older adult. To ensure downstream sample demultiplexing based on human leukocyte antigen (HLA) genotype was possible, paired media control and influenza antigen condition PBMC samples from the same subject were never combined in the same reaction. Multiplexed samples were loaded onto the Chromium Next GEM Single Cell 5’ assay using the Chromium Next GEM Single Cell 5’ HT v2 kit (10x Genomics, CA, USA), per the manufacturer’s protocol. The amplified cDNA libraries were sequenced on an Illumina NovaSeq 6000 S4 using a 2x150 bp paired-end chemistry kit (Illumina, CA, USA), targeting 40,000 reads per cell for the gene expression libraries and 10,000 reads per cell for the cell surface protein libraries.

### 10.8. Sequence alignment and sample demultiplexing

Raw sequencing reads were aligned to a KIR-modified GRCh38 human genome reference (*34*), which incorporates all possible KIR genes, using *Cell Ranger* v7.1.0 or v7.2.0 (10x Genomics, CA, USA) with default parameters. Note that there were no updates from *Cell Ranger* v7.1.0 to v7.2.0 that might affect alignment, filtering, barcode counting or unique molecular identifier (UMI) counting. Samples were demultiplexed using *Souporcell* v2.5 (*53*) with subject samples identified by comparing the HLA genotype obtained from genotyping with the *in silico* inferred HLA genotype from sequencing reads using *arcasHLA* v0.5.0 (*54*).

### 10.9. KIR transcript reassignment

Although alignment to a KIR-modified GRCh38 human genome reference improves detection of KIR transcripts, the high sequence homology among KIR genes may result in transcript mismapping to closely related loci (*34*). To assess the potential for mismapping in our data, we examined the nucleotide distance between KIR alleles in the cohort using MEGA X (*55*). KIR allele sequences spanning the first 150 nucleotide bases – regions preferentially captured by the Chromium Next GEM Single Cell 5’ assay and encompassing the highly homologous exons 1 and 2 – were extracted from the IPD-KIR Database (*51*). Phylogenetic trees were constructed using the Neighbor-Joining method (*56*), with evolutionary distances calculated based on nucleotide base differences, which correspond to mismatches considered in read alignment (Figure S3). For the alleles present in this cohort, we found a high potential for mismapping among KIR2DL1/L2/L3/S1/S2 and between KIR3DL1/S1. To account for this, we conservatively aggregated gene expression for KIR2DL1/L2/L3/S1/S2 and KIR3DL1/S1 (Figure S4; Table S8). Additionally, a minor fraction of KIR transcripts were misassigned to KIR2DL5 (median 0.2%, range 0.1-0.4%) and KIR2DS5 (median 0.35%, range 0.1-1.1%). However, based on genotype, these reads could be reassigned – KIR2DL5-derived transcripts to either KIR2DS3 or KIR2DS5, and KIR2DS5-derived transcripts to KIR2DS3 – as no subject co-expressed KIR2DS3 and KIR2DS5. To facilitate downstream analysis, we also generated a “*Total KIR*” measure representing the aggregated expression of all KIR transcripts per cell.

### 10.10. Quality control

Filtered gene expression and cell surface protein count matrices were imported into R v4.2.3 for additional quality control and downstream analyses using *Seurat* v4.3.0 (*57*). Quality control was performed for each single-cell reaction based on gene expression, as previously described (*58*). Briefly, dynamic thresholding of the count distribution was performed, which determined < 500 unique genes and < 1,000 total unique molecular identifiers (UMIs) as exclusion criteria for low quality cells. One reaction was found to have below the recommended cDNA input for sequencing due to low cell capture and therefore required an altered low quality cell threshold of < 250 unique genes and < 500 total UMIs. Moreover, cells with a mitochondrial gene content greater than three median absolute deviations above the median were similarly excluded, which equated to a median cut-off of 4.38% (range, 3.56-7.58%) mitochondrial gene content.

While doublets were previously identified in our companion paper (*19*) using *Souporcell* v2.5 (*53*), we performed confirmatory doublet detection using three additional methods: (1) Co-expression-based doublet scoring using *scds* v1.14.0 (*59*); (2) Binary classification-based doublet scoring using *scds* v1.14.0 (*59*); and (3) Iterative classification of real cells and artificial doublets using *scDblFinder* v1.15.4 (*60*). All three methods have been shown to have good all-around performance on PBMC datasets (*61*). Cells classified as singlets across as a minimum of three out of four methods were retained for downstream analysis.

### 10.11. NK cell extraction

NK cells were identified using a multi-step approach combining cluster- and cell-based methods.

#### Step 1: Integrated downsampling and clustering

To mitigate sequencing depth variability across batches, all samples were integrated by downsampling to ∼1,000 UMIs per cell, as previously described (*62*). The 1,000 UMI threshold was selected based on prior evidence demonstrating its sufficiency for cell type classification (*62, 63*). Downsampled data were normalized to Counts Per Million (CPM) and log-transformed. The top 5,000 variable features were selected, and dimensionality was reduced using principal component analysis (PCA; first 50 components) and Uniform Manifold Approximation and Projection (UMAP). In this approach, NK cells did not form a distinct cluster but were embedded within a larger T cell cluster. NK-like populations were manually extracted based on high expression of *Total KIR*, *FCGR3A*, *NCAM1*, *KLRK1*, *NKG7*, *GNLY,* and low expression of *CD14*, *LYZ*, *CD19*, *CD79A*, *CD3E*, *CD3D*, *CD4*, *CD8A*.

These NK-like populations comprised ∼122,000 cells.

#### Step 2: Default Seurat clustering per sample

To validate the NK cells identified in Step 1, an alternative clustering pipeline was applied using Seurat (*57*). Each sample was processed separately using Seurat’s default pipeline: normalization, selection of the top 2,000 variable features, scaling, PCA and UMAP. NK cell clusters were identified using the same marker criteria as in Step 1, yielding ∼61,000 total cells. Of these, 97% (∼59,000) overlapped with those identified in Step 1.

#### Step 3: Cell-based refinement

To refine the dataset, unique NK-like cells identified across Step 1 and Step 2 were combined and downsampled as in Step 1. Residual gene expression of *CD3D*, *CD3E* and *CD8A* suggested the presence of T cells, prompting a cell-based classification approach. *SingleR* v2.0.0 (*64*) was applied to annotate cells using three reference datasets from *celldex* v1.12.0: (1) Human Primary Cell Atlas (*65*); (2) Blueprint/ENCODE (*66, 67*); and (3) Database of Immune Cell Expression (*68*). Approximately 64,000 NK cells were consistently identified across all three references. Subsequent validation using *ScType* (*69*) confirmed 42,368 cells as NK cells, which served as the final NK cell dataset for downstream analysis.

### 10.12. Data integration, dimensionality reduction and cluster annotation

The final dataset, comprising 42,368 NK cells, underwent integration, dimensionality reduction and cluster annotation using the downsampling method described above (see NK cell extraction; Step 1). This method has been shown to improve integration by preserving biological variability while mitigating overcorrection (*62*). KIR, mitochondrial, and ribosomal genes were excluded from feature selection to prevent biases in clustering. Given that the dataset consists of a single cell type, 2,000 variable features were sufficient for dimensionality reduction. To determine the optimal clustering resolution, we used *clustree* v0.5.1 (*70*) to identify plateau points across resolutions ranging from 0.1 to 1.0. This analysis was guided by prior single-cell studies of NK cell heterogeneity in health human blood (*29*), which identified six NK cell populations under unstimulated conditions. Based on these insights, we selected a Leiden clustering resolution of 0.5, yielding eight clusters encompassing media control and influenza antigen conditions.

### 10.13. Lineage trajectory analysis

Lineage inference was performed using pseudotime trajectory analysis to assess cluster connectivity. The optimal pseudotime method was selected using *dyno* v0.1.2 (*71*), which evaluated 40 different trajectory inference methods. Among these, *slingshot* v2.6.0 (*72*) was identified as the most suitable method for our dataset based on its accuracy, scalability and stability. For the trajectory inference, *slingshot* was run using a mutual nearest neighbor-based distance metric, and the granularity parameter (*omega*) was included to identify outlier clusters.

## 11. Quantification and statistical analysis

All statistical analyses were conducted in R v4.2.3 running on Windows 10 x64. Group comparisons were performed using one of the following tests, as appropriate: repeated-measures ANOVA (*stats* v4.2.3), Wilcoxon signed-rank test (*stats* v4.2.3), Mann-Whitney U test (*stats* v4.2.3), or paired moderated t-test with false discovery rate (FDR) correction where applicable (*scuttle* v.1.8.4) (*73*). Statistical significance was defined as *p*-value (or FDR) < 0.05.

Differential expression analysis was performed using Seurat (*57*) with the DESeq2 pseudobulk method (*74*), aggregating gene expression at the subject and cluster level. Genes were considered differentially expressed if they met an adjusted *p*-value threshold of < 0.05 and an average log_2_ fold change (log_2_FC) > |2|.

Significant differences in subject characteristics were assessed using Fisher’s exact test (*stats* v4.2.3), with statistical significance defined as *p*-value < 0.05.

