## Supplementary Figures for "Adaptive-like NK cell responses to influenza correlate with humoral immunity and are influenced by age and sex"

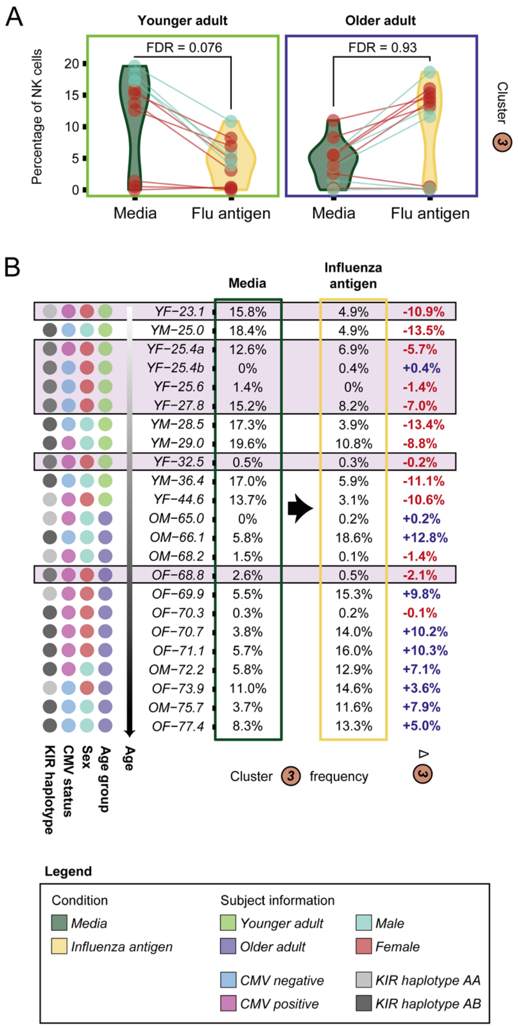


**Figure S1. Cluster 3 shows opposing trends with age but lacks statistical significance. A)** Change in cluster 3 frequency between media control and influenza antigen conditions, shown separately for each age group, as in Figure 2B. **B)** Frequency of cluster 3 cells by subject, stratified by condition. Subjects are ordered by age, with sex, cytomegalovirus (CMV) status and killer immunoglobulin-like receptor (KIR) haplotype indicated. An increased frequency following influenza antigen stimulation is shown in blue, whereas a decreased frequency is indicated in red. Subjects with major increases (>50%) in cell frequency for clusters 4 and/or 5 in the presence of influenza antigen (as identified in Figure 4A) are highlighted in purple. No clear associations, aside from age, were observed in the pattern of cluster 3 frequency changes in the presence of influenza antigen.

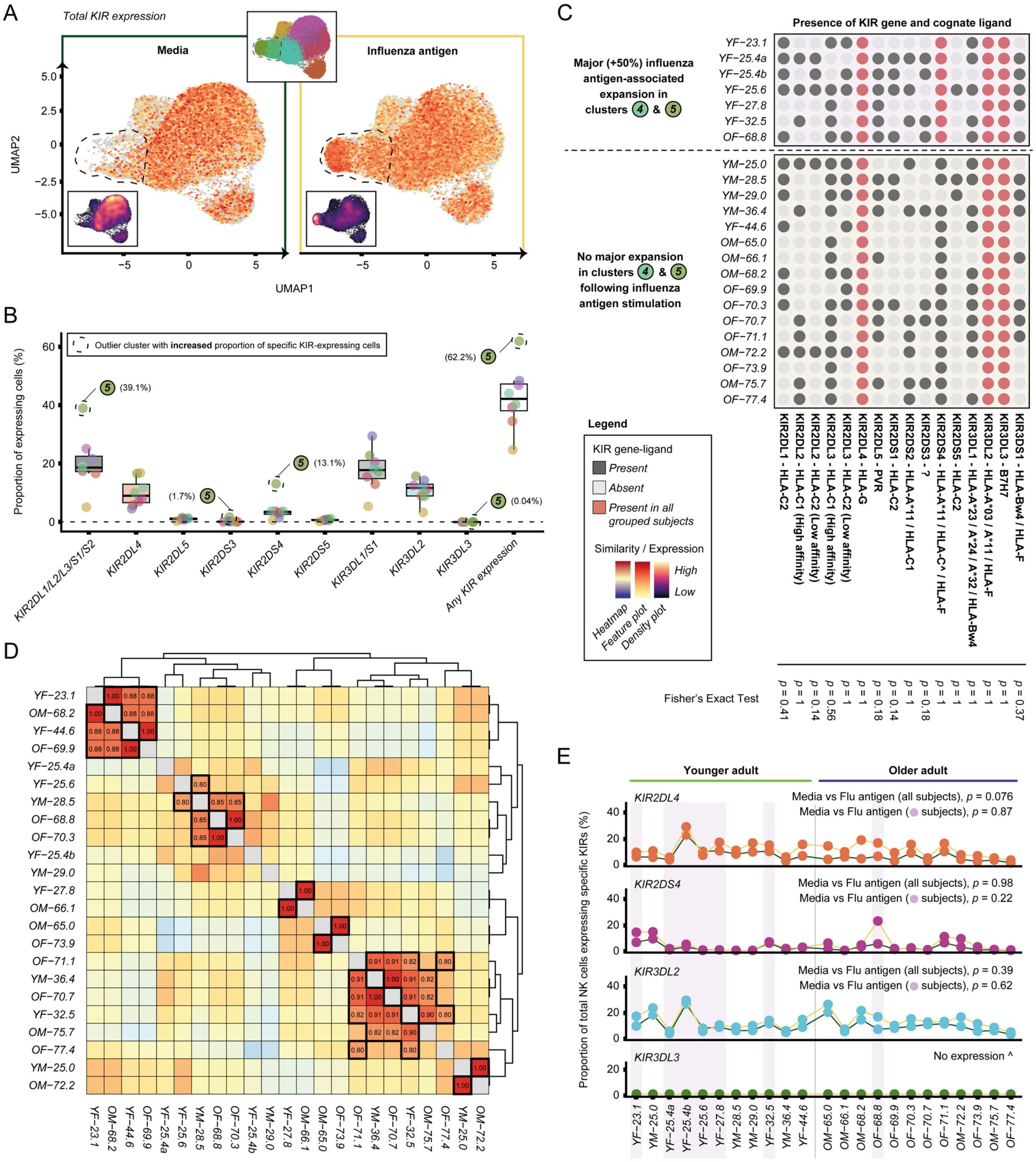


**Figure S2. KIR gene-ligand repertoire does not associate with influenza antigen-associated NK cell reactivity. A)** Feature plot and corresponding density plot showing total KIR gene expression, stratified by condition. Dashed region indicates clusters 4 and 5. **B)** Boxplot showing the frequency of cells expressing specific KIR genes by cluster. Boxes represent the median and interquartile range (IQR). Whiskers extend to frequencies within 1.5 x IQR from the first and third quartiles. Outlier clusters having increased frequency of cells expressing specific KIR genes are indicated. **C)** Presence or absence of KIR genes and their cognate ligand(s) by subject. A dark-filled circle indicates the presence of both the KIR gene and at least one cognate KIR ligand, while an empty circle indicates that either the KIR gene or its cognate ligand is absent. A red-filled circle indicates a KIR gene-ligand combination that is present in all grouped subjects. Fisher's Exact Test shows that no KIR gene-ligand combination was significantly enriched among the seven grouped subjects who exhibited major increases in clusters 4/5 in the presence of influenza antigen. **D)** Heatmap of the KIR gene-ligand repertoire by subject, based on the Jaccard distance calculated on the number of present KIR gene-ligand combinations. Subjects with a high degree of KIR gene-ligand repertoire similarity are indicated. **E)** Dot plot showing the frequency of NK cells expressing specific KIR genes by subject and condition. KIR genes assessed are those present in all grouped subjects (red-filled circles in (C)). Comparisons between media and influenza antigen conditions were made between all subjects ($n$ = 23) and the subset of subjects with major increases in clusters 4/5 ($n$ = 7) using a paired moderated t-test. The frequency of KIR3DL3-expressing cells was zero in 22/23 subjects and detected in only one cell in the remaining subject.

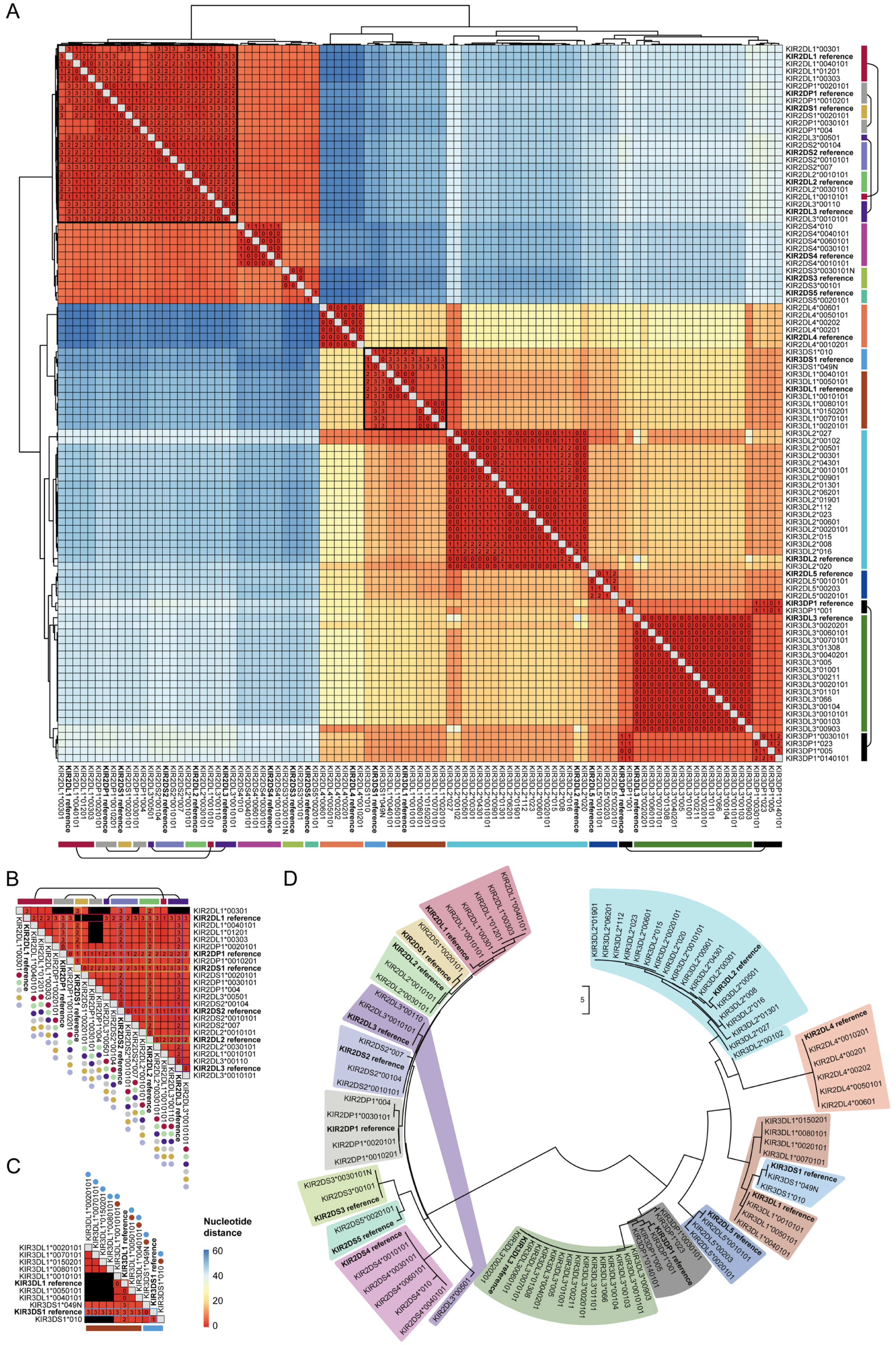
A
B
C
D
Nucleotide distance
60
40
20
0

**Figure S3. KIR allele sequence homology across the cohort. A)** Heatmap of nucleotide distance between KIR alleles for the examined subjects and reference KIR genes used in the KIR-modified GRCh38 genome. **B & C)** Detailed views of regions with high sequence homology. **D)** Phylogenetic analysis of KIR alleles present across examined subjects. The Neighbor-Joining method was used, and the tree is drawn to scale, with branch lengths representing the number of nucleotide base differences. KIR allele transcript sequences spanning the first 150 nucleotide bases (relevant region for the 5’ single-cell chemistry) were analyzed.

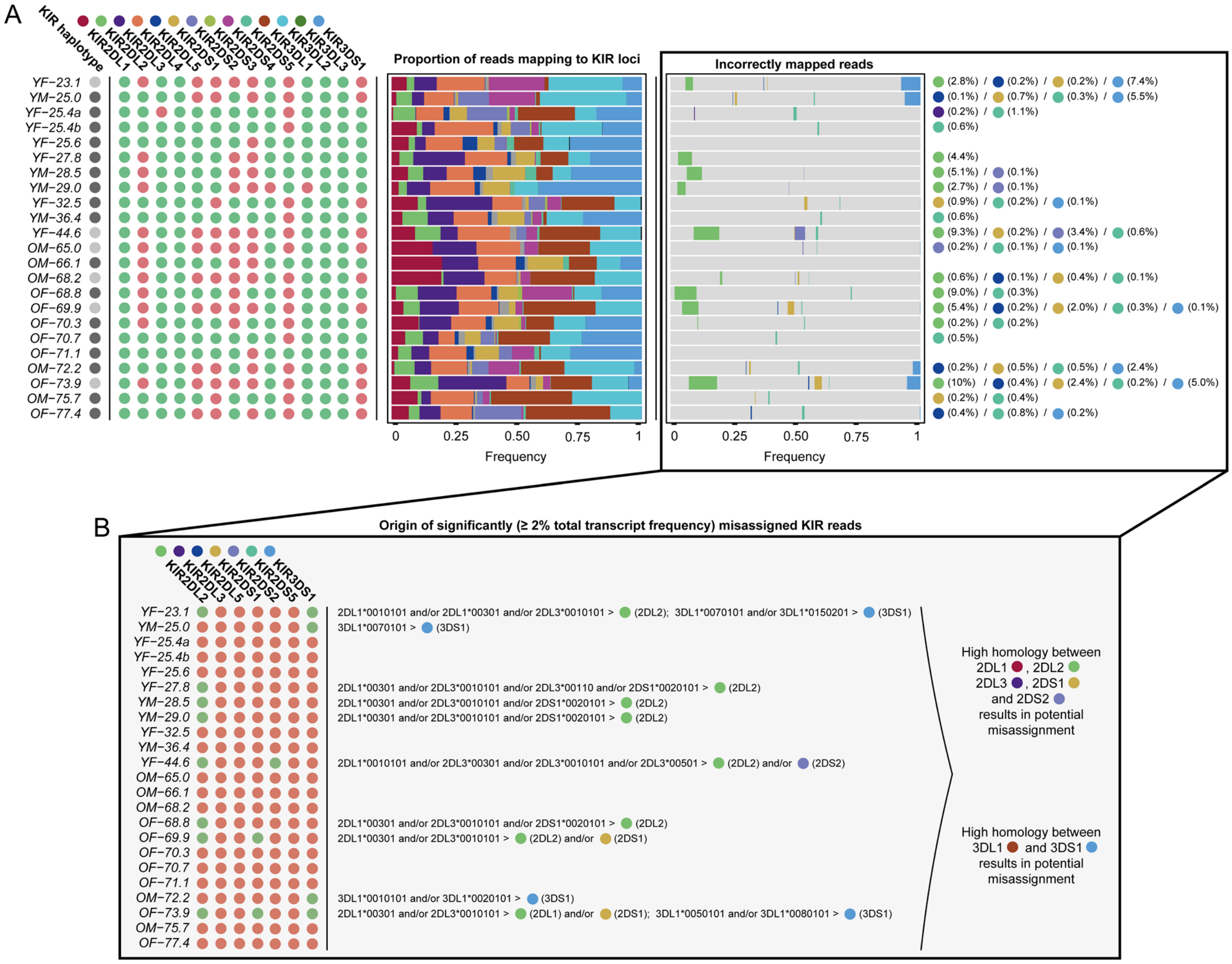


**Figure S4. Identification of mismapped KIR reads. A)** KIR genotyping, proportion of KIR reads and genotyping-informed identification of mismapped KIR reads across the examined subjects. **B)** Sequence homology, as determined in Figure S3, was used to trace the origin of high-frequency (≥ 2%) misassigned KIR reads. As predicted in Figure S3, significant misassignment was attributed to highly homologous KIR2DL1/L2/L3/S1/S2 and KIR3DL1/S1 transcripts at the 150 nucleotide transcript capture length.
